# Non-linear randomized Haseman-Elston regression for estimation of gene-environment heritability

**DOI:** 10.1101/2020.05.18.098459

**Authors:** Matthew Kerin, Jonathan Marchini

## Abstract

Gene-environment (GxE) interactions are one of the least studied aspects of the genetic architecture of human traits and diseases. The environment of an individual is inherently high dimensional, evolves through time and can be expensive and time consuming to measure. The UK Biobank study, with all 500,000 participants having undergone an extensive baseline questionnaire, represents a unique opportunity to assess GxE heritability for many traits and diseases in a well powered setting. We have developed a non-linear randomized Haseman-Elston (RHE) regression method applicable when many environmental variables have been measured on each individual. The method (GPLEMMA) simultaneously estimates a linear environmental score (ES) and its GxE heritability. We compare the method via simulation to a whole-genome regression approach (LEMMA) for estimating GxE heritability. We show that GPLEMMA is computationally efficient and produces results highly correlated with those from LEMMA when applied to simulated data and real data from the UK Biobank.

## Introduction

The advent of genome-wide association studies^1^ has catalyzed a huge number of discoveries linking genetic markers to many human complex diseases and traits. For the most part these discoveries have involved common variants that confer relatively small amounts of risk and only account for a small proportion of the phenotypic variance of a trait^2^. This has led to a surge of interest in methods and applications that measure the joint contribution to phenotypic variance of all measured variants throughout the genome (SNP heritability), and in testing individual variants within this framework. Most notably the seminal paper of Yang et al. (2010), who used a linear mixed model (LMM) to show that the majority of missing heritability for height could be explained by genetic variation by common SNPs^3^. When testing variants for association these LMMs can reduce false positive associations due to population structure, and improve power by implicitly conditioning on other loci across the genome^4–6^. These methods model the unobserved polygenic contribution as a multivariate Gaussian with covariance structure proportional to a genetic relationship matrix (GRM) ^7–9^. This approach is mathematically equivalent to a whole genome regression (WGR) model with a Gaussian prior over SNP effects ^4^.

Subsequent research has shown that the simplest LMMs make assumptions about the relationship between minor allele frequency (MAF), linkage disequilibrium (LD) and trait architecture that may not hold up in practice^10,11^ and generalisations have been proposed that stratify variance into different components by MAF and LD^10, 12, 13^. Other flexible approaches have been proposed in both the animal breeding^14, 15^ and human literature^16–18^ to allow different prior distributions that better capture SNPs of small and large effects. For example, a mixture of Gaussians (MoG) prior can increase power to detect associated loci in some (but not all) complex traits^6, 17^. Other methods have been proposed that estimate heritability only from summary statistics and LD reference panels ^19, 20^. Heritability can also be estimated using Haseman-Elston regression ^21^ and has recently been extended using a randomised approach ^22^ that has 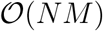 computational complexity and works for multiple variance components ^23^. Other recent work has shown that LMM approaches such as these are not able to disentangle direct and indirect genetic effects, the balance of which will vary depending on the trait being studied. ^24^.

There has been less exploration of methods for estimating heritability that account for gene-environment interactions. One interesting approach has proposed using spatial location as a surrogate for environment^25^ using a three component LMM - one based on genomic variants, one based on measured spatial location as a proxy for environmental effects, and a gene-environment component, modeled as the Hadamard product of the genomic and spatial covariance matrices. Other authors have used this method to account for gene-gene interactions ^26, 27^.

Modelling gene-environment interactions when many different environmental variables are measured is a more challenging problem. If several environmental variables drive interactions at individual loci, or if an unobserved environment that drives interactions is better reflected by a combination of observed environments, it can make sense to include all variables in a joint model. StructLMM^28^ focuses on detecting GxE interactions at individual markers, and models the environmental similarity between individuals (over multiple environments) as a random effect, and then tests each SNP independently for GxE interactions. However this approach does not model the genome wide contribution of all the markers, which is often a major component of phenotypic variance.

We recently proposed a whole genome regression approach called LEMMA applicable to large human datasets such as UK Biobank, where many potential environmental variables are available^29^. The LEMMA regression model includes main effects of each genotyped SNP across the genome, and also interactions of each SNP with a environmental score (ES), that is a linear combination of the environmental variables. The ES is estimated as part of the method. The model uses mixture of Gaussian (MoG) priors on main and GxE SNP effects, that allow for a range of different genetic architectures from polygenic to sparse genetic effects^16–18^. The ES can be readily interpreted and its main use is to test for GxE interactions one variant at a time, typically at a larger set of imputed SNPs in the dataset. However, the ES can also be used to estimate the proportion of phenotypic variability that is explained by GxE interactions (SNP GxE heritability), using a two component randomised Haseman-Elston (RHE) regression ^23^.

The main contribution of this paper is to combine the estimation of the LEMMA ES into a standalone RHE framework. This results in a non-linear optimization problem that we solve using the Levenburg-Marquardt (LM) algorithm. The method implicitly assumes a Gaussian prior on main effect and GxE effect sizes. We also propose a separate RHE method that estimates the independent GxE contribution of each measured environmental variable. We set out the differences between these two models and present a simulation study to compare them to LEMMA. We also apply the method to UK Biobank data and show that GPLEMMA produces estimates very close to LEMMA. Software implementing the GPLEMMA algorithm in C++ is available at https://jmarchini.org/gplemma/.

## Methods

### Modeling SNP Heritability

The simplest model for estimating SNP heritability has the form

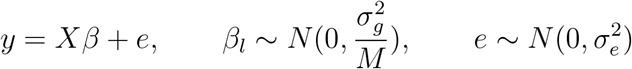

where *y* is a continuous phenotype, *X* is an *N* × *M* matrix of genotypes that has been normalised to have column mean zero and column variance one, and *β* is an M-vector of SNP effect sizes. Integrating out *β* leads to the variance component model

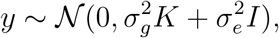

where 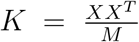 is known as the genomic relationship matrix (GRM)^3^. Estimating the two parameters in this model *σ_g_* and *σ_e_* leads to an estimate of SNP heritability of 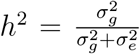. This is commonly referred to in the literature as the single component model. Subsequent research has shown that the single component model makes assumptions about the relationship between minor allele frequency (MAF), linkage disequilibrium (LD) and trait architecture that may not hold up in practice^10, 11^. There have been many follow up methods, including; generalizations that stratify variance into different components by MAF and LD ^13^, approaches that assign different weights for the GRM^10, 12^, methods that replace the Gaussian prior on *β* with a spike and slab on SNP effect sizes ^30^ and methods that estimate heritability only from summary statistics and LD reference panels ^20, 31^.

### Haseman-Elston (HE) regression

An alternative method used to compute heritability is known as HE-regression^21^. HE-regression is a method of moments (MoM) estimator that optimizes variance components 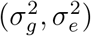 in order to minimise the squared difference between the observed and expected trait covariances. The MoM estimator 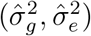 can be obtained by solving the minimization

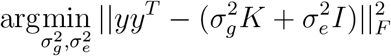

or equivalently by solving the linear regression problem

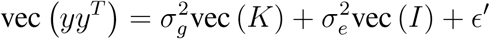

where vec(*A*) is the vectorization operator that transforms an *N × M* matrix into an *NM*-vector.

In matrix format, both of these forms correspond to solving the following linear system

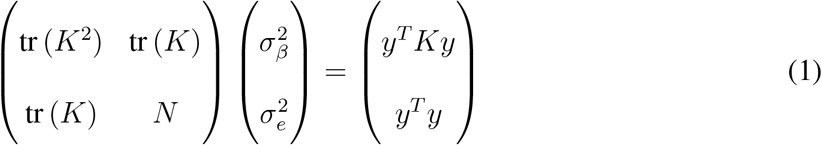

HE-regression methods are widely acknowledged to be more computationally efficient^22, 32, 33^ and do not require any assumptions on the phenotype distribution beyond the covariance structure ^32^ (in contrast to maximum-likelihood estimators). However, HE-regression based estimates typically have higher variance ^33^, thus implying that they have less power.

Recent method developments^22, 23^ have shown that a randomized HE-regression (RHE) approach can be used to compute efficiently on genetic datasets with hundreds of thousands of samples.

Wu et al. (2018) observed that Equation (1) can be solved efficiently without ever having to explicitly compute the kinship matrix *K* by using Hutchinson’s estimator ^34^, which states that tr 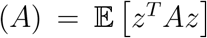 for any matrix where *z* is a random vector with mean zero and covariance given by the identity matrix. The proposed method involves approximating tr (*K*) and tr (*K*^2^) using only matrix vector multiplications with the genotype matrix *X*, to compute the following expressions

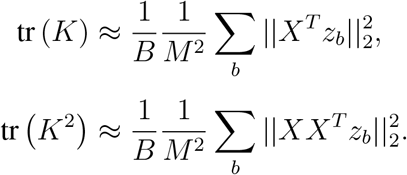

Thus an approximate solution can be obtained in 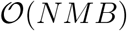 time, where *B* denotes a relatively small number of random samples. Subsequent work by extended this approach to a multiple component model ^23^

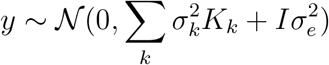

With parameter estimates obtained as solution to the linear system given by

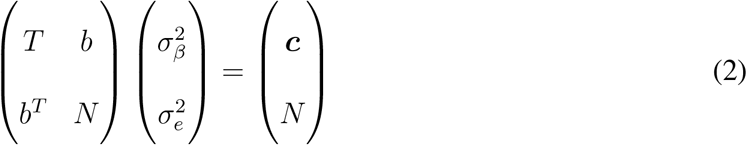

where *T_kl_* = tr (*K_k_K_l_*), *b_k_* = tr (*K_k_*) and *c_k_* = *y^T^K_k_y*. Finally both papers show how to efficiently control for covariates by projecting them out of of all terms in the system of equations. Thus with covariates included the multiple component model becomes

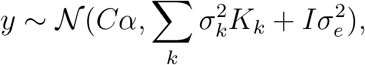

and terms in the subsequent linear system are given by *T_kl_* = tr (*WK_k_WK_l_W*), *b_k_* = tr (*WK_k_W*) and *c_k_* = *y^T^WK_k_Wy*, where *W* = *I_N_* − *C^T^*(*C^T^C*)^-1^*C*.

### Modeling GxE heritability

We introduce two extensions of the RHE framework for modelling GxE interactions with multiple environmental variables. In both models we let *E* be an *N* × *L* matrix of environmental variables, C is an N × *D* matrix of covariates, each with columns normalised to have mean zero and variance one.

### MEMMA

The first model assumes that each environmental variable interacts independently with the genome

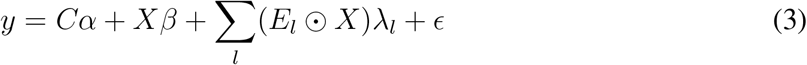

where 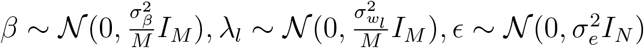 and *E_l_*ʘ*X* denotes the element-wise product of *E_l_* with each column of *X*. Integrating out *β* and *λ* leads to the variance component model

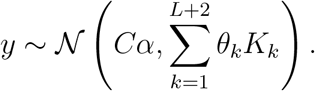

where 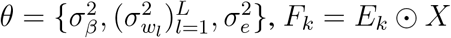 and

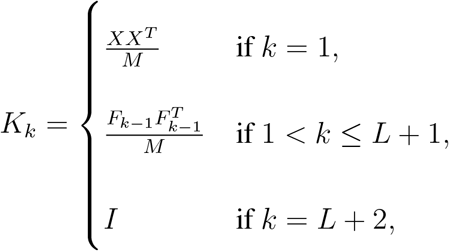

Fitting the variance components is done analytically by solving the system of equations *Tθ* = *c* where *T_kl_* = tr (*WK_k_WK_l_W*), *c_k_* = *y^T^WK_k_Wy* and *W* = *I_N_* − *C*(*C^T^C*)^-1^*C^T^*. As shown in the original RHE method^22, 23^, Hutchinson’s estimator can be used to efficiently estimate *T_kl_*. To do this our software streams SNP markers from a file and computes *y^T^WXX^T^Wy* and the following *N*-vectors

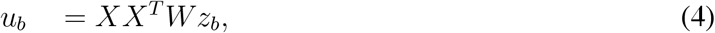

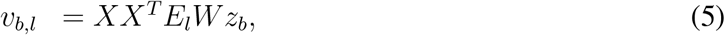

where *z_b_* ~ *N*(0, *I_N_*)for 1 ≤ *b* ≤ *B* are random *N*-vectors. Then

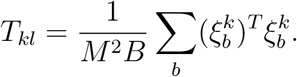

where 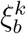 is defined as

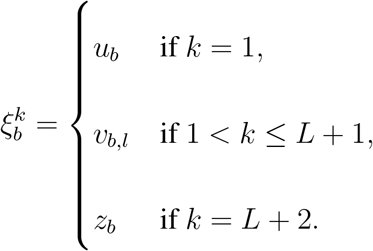

Finally, the variance components are converted to heritability estimates using the following formula

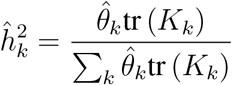

We call this approach MEMMA (Multiple Environment Mixed Model Analysis). MEMMA costs 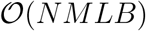 in compute and 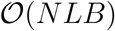 in RAM.

### GPLEMMA

The second model involves the estimating a linear combination of environments, or environmetal score (ES), that interacts with the genome. The model is given by

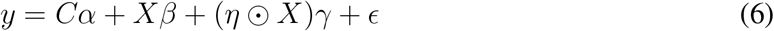

where *η = E_w_* is the linear environmental score (ES), 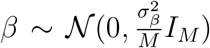 and 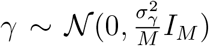. This is the same model used by LEMMA^29^ except the mixture of Gaussians priors on SNP effects (*β* and *γ*) have been replaced with Gaussian priors. For this reason we call this approach GPLEMMA (Gaussian Prior Linear Environment Mixed Model Analysis). Integrating out the SNP effects yields the model

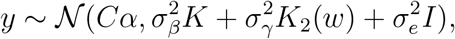

where 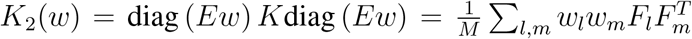 and *F_l_* = *E_l_* ʘ *X*. Minimizing the squared loss between the expected and observed covariance is equivalent to the following regression problem

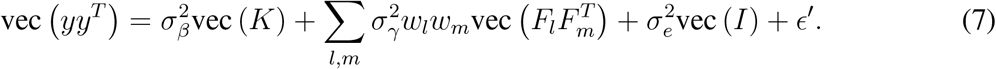

In this format it is clear that optimising 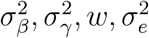 is a non-linear regression problem. Further, including a parameter for 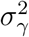 is no longer necessary. From here on we set 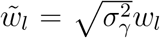 and drop the 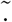 parameterisation without loss of generality.

### Levenburg-Marquardt algorithm

We use the Levenburg-Marquardt (LM) algorithm^35^, which is commonly used for non-linear least squares problems. The algorithm effectively interpolates between the Gauss-Newton algorithm and the method of steepest gradient descent, by use of an adaptive damping parameter. In this manner, it is more robust than the straight forward Gauss-Newton algorithm but should have faster convergence than a gradient descent approach.

Without loss of generality, consider the model

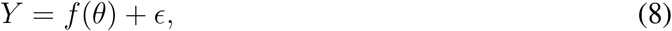

where *f*(*θ*) is a function that is non-linear in the parameters *θ*. Given a starting point *θ*_0_, LM proposes a new point *θ*_new_ = *θ*_0_ + *δ* by solving the normal equations

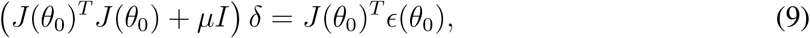

where 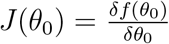 and *ϵ*(*θ*_0_) = *Y* − *f*(*θ*_0_) are respectively the Jacobian and the residual vector evaluated at *θ*_0_.

If *θ*new has lower squared error than *θ*_0_, then the step is accepted and the adaptive damping parameter *μ* is reduced. Otherwise, *μ* is increased and a new step *δ* is proposed. For small values of *μ* Equation (9) approximates the quadratic step appropriate for a fully linear problem, whereas for large values of *μ* Equation (9) behaves more like steepest gradient descent. This allows the algorithm to defensively navigate regions of the parameter space where the model is highly non-linear. If *θ* + *δ* reduces the squared error, then the step is accepted and *μ* is reduced, otherwise *μ* is increased and a new step *δ* is proposed.

In summary the LM algorithm requires computation of the matrices *J*(*θ*)^*T*^*J*(*θ*), *J*(*θ*)^*T*^*ϵ*(*θ*) at each step, as well as the squared error (which we define as *S*(*θ*)). We now give statements of the equations used to compute each of these values, and show that each iteration can be performed in 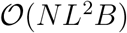 time.

We apply the LM algorithm with 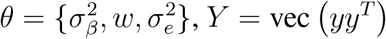 and

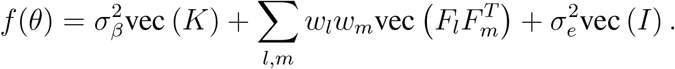

Several quantities can be pre-calculated and re-used in the LM algorithm. The N-vectors *u_b_*, *v_b,l_*, and *y^T^WXX^T^Wy* are needed and have been defined above. In addition, GPLEMMA also benefits from the pre-calculation of

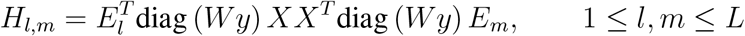

which can also be computed as genotypes are streamed from file.

Let (*J^T^ J*)*_θ_i__,_θ_j__* denote the entry of the *J^T^J* that corresponds to 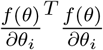 for 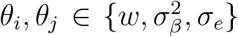 and define the *N*-vector *v_b_*(*w*) = ∑*_l_w_l_v_bl_*. Then the (*L* + 2) × (*L* + 2) matrix *J*(*θ*)^T^*J*(*θ*) is given by

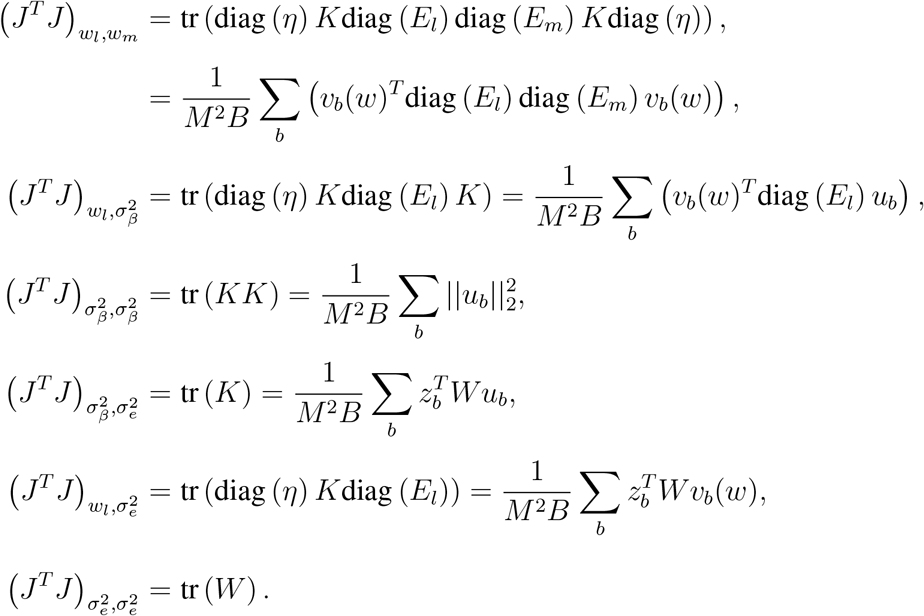

*J*(*θ*)^*T*^*ϵ*(*θ*) is given by

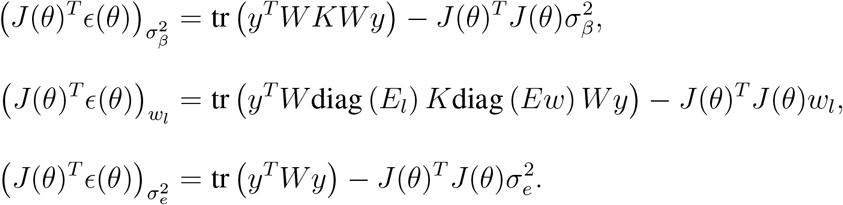

where

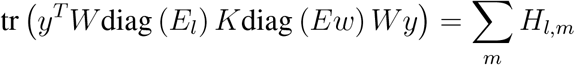

Finally the squared error, which we define as *S*(*θ*), is given by

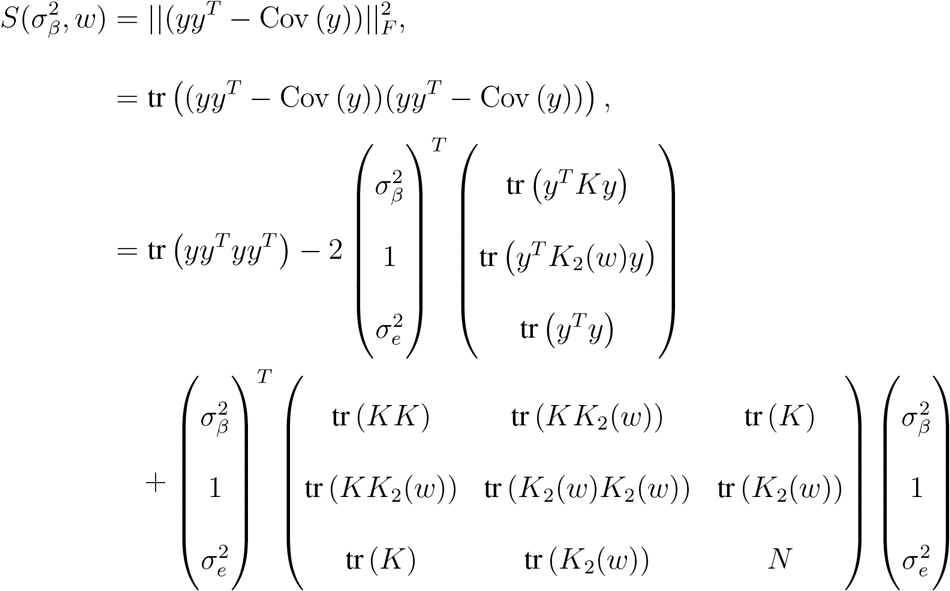

where

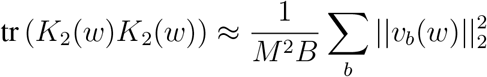

The initial preprocessing step has costs 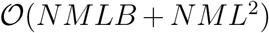 in compute and 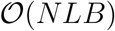 in RAM. The remaining algorithm does not require much RAM in addition to that required in the preprocessing step, so also costs 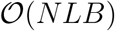 in RAM. Construction of the summary variable *v_b_*(*w*) = ∑*_i_w_l_v_b,l_* costs 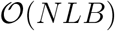 in compute. Each iteration of the LM algorithm costs 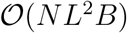.

It is possible to parallelise GPLEMMA using OpenMPI by partitioning samples across cores, in a similar manner to that used by LEMMA ^29^. Given that evaluating the objective function 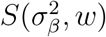 is characterised by BLAS level 1 array operations, a distributed algorithm using OpenMPI should have superior runtime versus an the same algorithm using OpenMP as well as providing RAM limited only by the size of a researchers compute cluster.

We perform 10 repeats of the LM algorithm with different initialisations, and keep results from the solution with lowest squared error 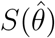. Each run is initialised with a vector of interaction weights 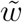, where each entry set to 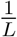 and a small amount of Gaussian noise is added.

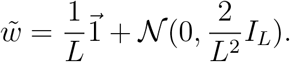

To transform the initial weights vector 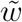 to the initial parameters *θ*_0_ we let 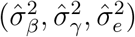 be solutions to

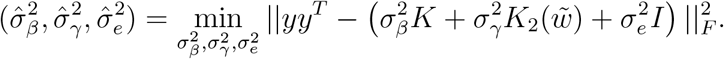

The GPLEMMA algorithm is then initialized with 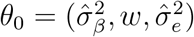 where 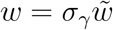.

### Relationship between MEMMA and GPLEMMA

Comparing Equation (3) with Equation (6), suggests that the GPLEMMA model can be expressed at the MEMMA model with the added constraint that

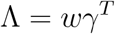

where Λ = {*λ*_1_,..., *λ_M_*} is the *L × M* matrix of GxE effect sizes in MEMMA for the *L* environments and *M* SNPs.

We can expect the two models to give similar heritability estimates, under the simplifying assumptions that GxE interactions do occur with a single linear combination of the environments and that the set of random variables 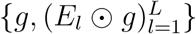 is mutually independent. Let 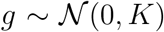 and 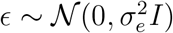. Then connection between the two models is revealed by observing

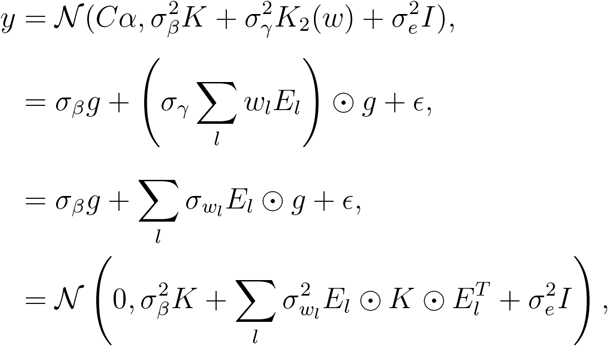

where 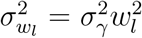. Thus we should expect both models to have the same estimate for the proportion of variance explained by GxE interaction effects.

Even in that case that MEMMA and GPLEMMA have the same expected heritability estimate, there are still some differences between the two. GPLEMMA is a constrained model, so the variance of its heritabiity estimates may be smaller. Further, although 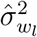 is proportional to the square of the weights used to construct the ES the sign of the interaction weight *w_l_* has been lost. Thus it is not possible to reconstruct an ES for use in single SNP testing using MEMMA.

## Results

### Simulated data

We carried out a simulation study to assess the relative properties of MEMMA and GPLEMMA. In addition, we compared to running the whole genome regression model in LEMMA, which estimates an ES and then uses it to estimate the GxE heritability.

The simulations use real data subsampled from genotyped SNPs in the UK Biobank^36^, drawing SNPs from all 22 chromosomes in proportion to chromosome length and using unrelated samples of mixed ancestry (N = 25k; 12500 white British, 7500 Irish and 5000 white European, *N* = 50k; 29567 white British, 7500 Irish and 12568white European, *N* = 100k; 79567 white British, 7500 Irish and 12568 white European; using self-reported ancestry in field *f.21000.0.0*). All samples were genotyped using the UKBB genotype chip and were included in the internal principal component analysis performed by the UK Biobank. Environmental variables were simulated from a standard Gaussian distribution.

Phenotypes for the baseline simulations were all simulated according to the LEMMA model ^29^. Let *N* be the number of individuals, *M* the total number of SNPs, *Mg* the number of causal main effect SNPs, *M_GxE_* the number of SNPs with GxE effects, *L* the total number of environmental variables, L^active^ the number of ’active’ environments with non-zero contribution to the ES vector *w*, and 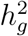 and 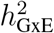 the herirtability of main effects and GxE effects. The model used to simulate data is

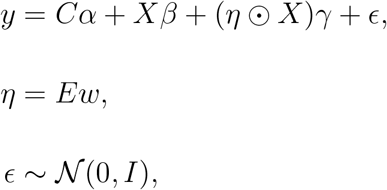

where *X* represents the *N* × *M* genotype matrix after columns have been standardised to have mean zero and variance one, *C* is the first principle component of the genotype matrix and *E* is the *N × L* matrix of environmental variables. In all simulations *α* was set such that *Cα* explained one percent of trait variance.

Non-zero elements of the interaction weight vector *w* were drawn from a decreasing sequence

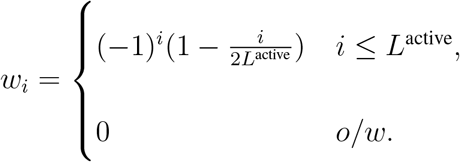

The effect size parameters *β* and *γ* were simulated from a spike and slab prior such that the number of non-zero elements was governed by *M_g_* and *M_GxE_* for main and interaction effects respectively. Non-zero elements were drawn from a standard Gaussian, and then subsequently rescaled to ensure that the heritability given by main and interaction effects was 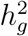 and 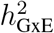 respectively. We chose a set of baseline parameter choices: *N* = 25*K*; *M =* 100K; *L* = 30; L^active^ = 6, *M_g_ =* 2500; M_GxE_ = 1250; 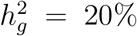; 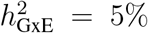, and then varied one parameter at a time to examine the effects of sample size, number of environments, number of non-zero SNP effects and GxE heritability. In addition, we investigated performance using a larger baseline simulation with *N* = 100K samples and *M* = 300K variants. The first genetic principal component was provided as a covariate to all methods.

Figure 1 compares estimates of the percentage variance explained (PVE) by GxE effects from all three methods. In general, all methods had upwards bias that decreased with sample size and increased with the number of environments. While heritability estimates from LEMMA and GPLEMMA appeared quite similar, estimates from MEMMA had much higher variance and also appeared to have higher upwards bias as the total number of environments increase. All the methods exhibited less bias in the larger simulations with *N* = 100K samples and *M* = 300K variants (Figure 1 (e-g)).

**Figure 1:**
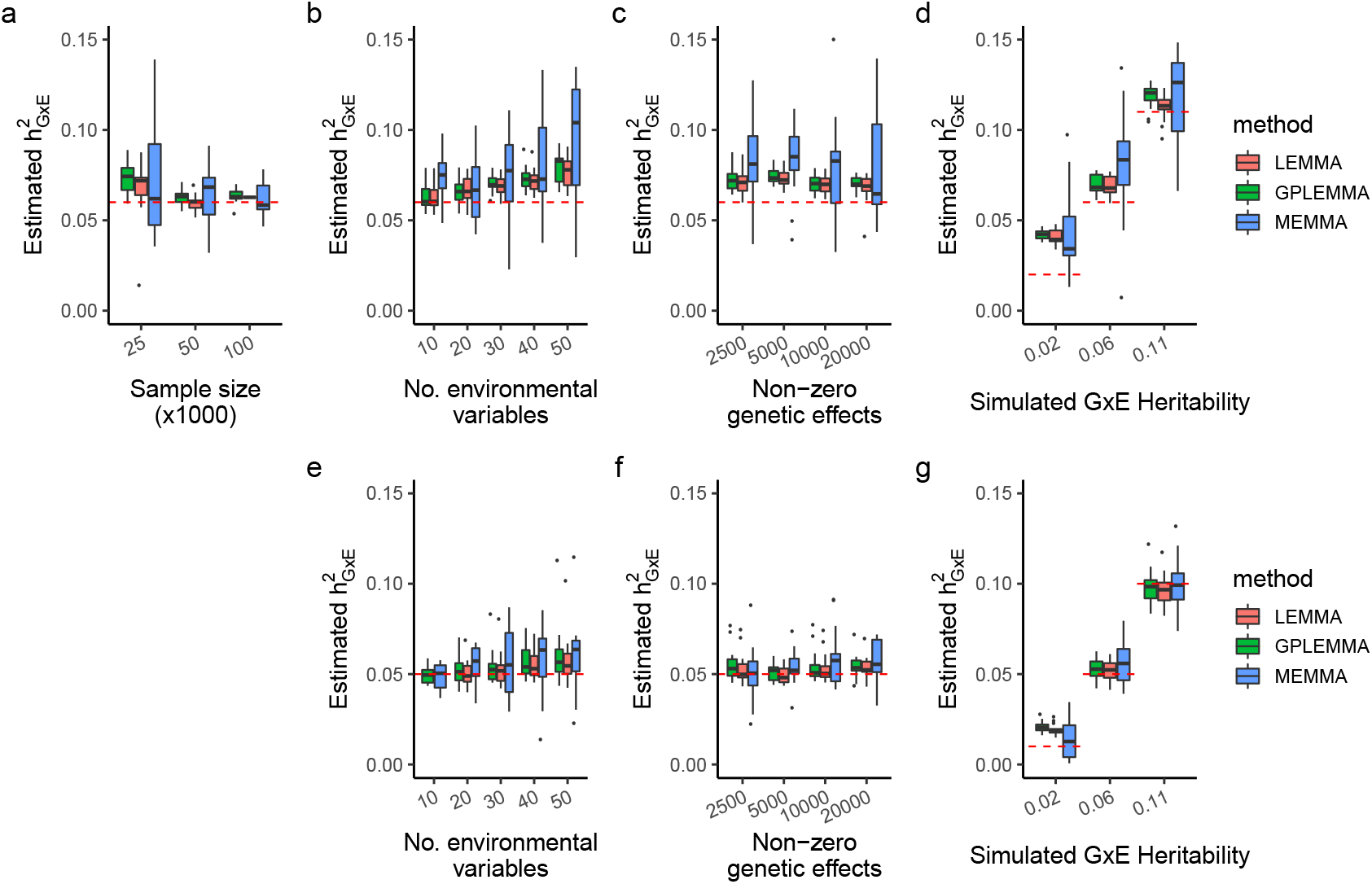
PVE estimation. Estimates of the proportion of variance explained by GxE effects by LEMMA, MEMMA and GPLEMMA on baseline simulations with using *N* = 25*K* samples and *M* = 100K variants, whilst varying sample size (a), the number of environments (b), the number of non-zero SNP effects (c) and GxE heritability (d). Panels (e-g) shows results of simulations with *N* = 100K samples and *M* = 300K variants, whilst varying the number of environments (e), the number of non-zero SNP effects (f) and GxE heritability (g).

Figure 2 compares the absolute correlation between the simulated ES and the ES inferred by LEMMA and GPLEMMA. Models like MEMMA do not provide an estimate of the ES. In general, the estimated ES from GPLEMMA had slightly lower absolute correlation with the true ES than the estimated ES from LEMMA, likely due to the data having been simulated from the LEMMA model, with sparse main and GxE SNP effects, whereas the GPLEMMA model assumes a polygenic or infinitesimal model. In large sample sizes (N = 100k), both methods achieve a correlation of over 0.98 with the simulated ES.

**Figure 2:**
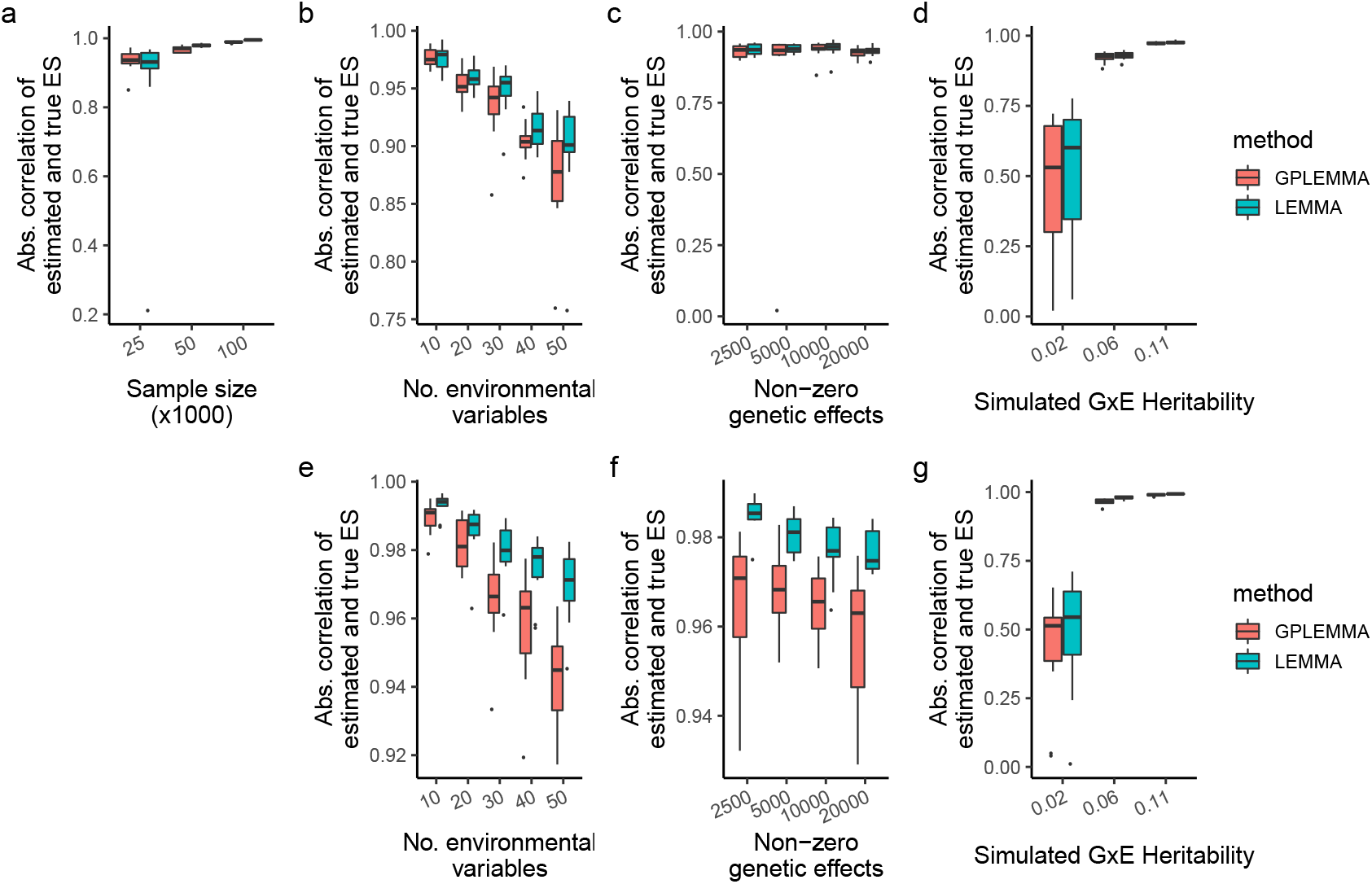
Comparison on baseline simulations. Absolute correlation between the true ES and the ES inferred by LEMMA and GPLEMMA whilst varying the number of environments, the number of active environments, the number of non-zero SNP effects and GxE heritability. The top row contains simulations using *N* = 25K samples and *M* = 100K variants, the bottom row containhs simulations using *N* = 100K samples and *M* = 300K variants. Results from 15 repeats shown.

Figure 3 compares MEMMA, GPLEMMA and LEMMA in a simulation where the functional form of a heritable environmental variable was misspecified (or more specifically; the phenotype depended on the squared effect of a heritable environment). All methods were first tested without any attempt to control for model misspecification, and second using a preprocessing strategy where each environment was tested independently for squared effects on the phenotype and any squared effects with p-value < 0.01/L were included as covariates. These are referred to as (*-SQE*) and (*+SQE*) respectively in the figures. Using the (*-SQE*) strategy, all methods showed upwards bias in estimates of GxE heritability that increased with the strength of the squared effect on the phenotype (Figure 3b). Model misspecification also caused bias in the ES of both GPLEMMA and LEMMA, however bias in the ES from GPLEMMA appeared to be much worse (Figure 3a). Using the (*+SQE*) strategy, all GxE heritability estimates were unbiased, consistent with earlier simulation results.

**Figure 3:**
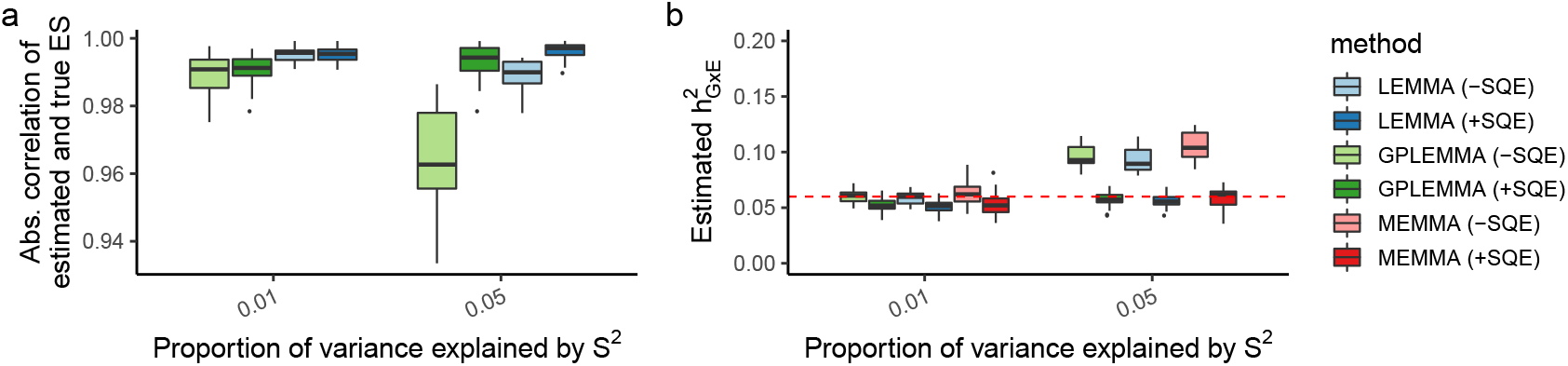
Comparison on simulations with a misspecified heritable environment. Estimated proportion of trait variance explained by GxE effects is shown on the left, absolute correlation between the inferred ES and the true ES shown in the right. Results shown using LEMMA, MEMMA and GPLEMMA. Phenotypes simulated with a squared effect from a heritable confounder (see **??**). Results from 20 repeats shown. Abbreviations; (-SQE), no attempt to control for squared effects; (+SQE), squared effects with *p* < 0.01 (Bonferroni correction for multiple envs) included as covariates

Figure 4 displays simulation results on the computational complexity of GPLEMMA. Figures 4a and 4b show that GPLEMMA achieved perfect strong scaling^1^ on the range of cores tested. This suggests that GPLEMMA has superior scalability to LEMMA, as for LEMMA the speedup due to increased cores began to decay after the number of samples per core dropped below 3000 ^29^.

**Figure 4:**
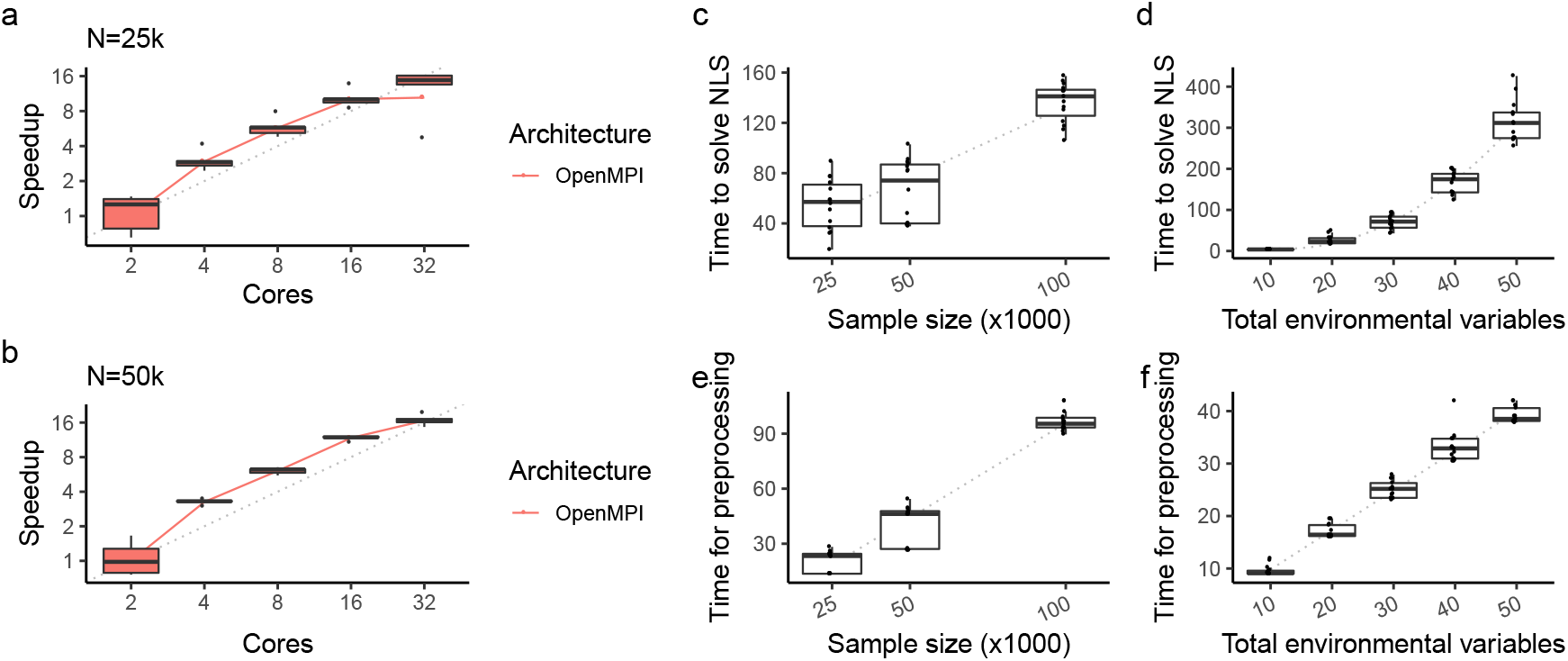
Computational complexity of GPLEMMA in simulation. Strong scaling of GPLEMMA using OpenMPI to parrallelise across cores with (a) *N* = 25*k* samples and (b) *N* = 50k samples. Comparison of the runtime of the Levenburg-Marquardt algorithm (c, d) and runtime of the preprocessing step (e, f). By default each run used; four cores, *N* = 25*k* samples, *L* = 30 environments and 10 random starts of the Levenburg-Marquardt algorithm. Results from 15 repeats shown.

Time to compute the preprocessing step and solve the non-linear least squared problem are shown in Figures 4c to 4f, while the number of environments and sample size were varied. As expected, the preprocessing step appeared to be linear in both the number of environments and sample size. Time to solve the non-linear least squares problem appeared to be quadratic in the number of environments and approximately linear in sample size *N*. As a single LM iteration should have complexity 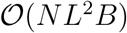, this suggests that the number of iterations required for convergence of the LM algorithm was independent of sample size and the number of environments (at least for the range of values tested).

Finally, to give a direct comparison between LEMMA and GPLEMMA, we ran each method on simulated data with *N* = 100*k* samples, *M* = 100*k* SNPs and *L* = 30 environmental variables using 4 cores for each run. Over 20 repeats, LEMMA took an average of 648 minutes to run whereas GPLEMMA took an average of 233 minutes.

### Analysis of UK Biobank data

To compare GPLEMMA and LEMMA on real data we ran both methods on body mass index (log BMI), systolic blood pressure (SBP), diastolic blood pressure (DBP) and pulse pressure (PP) measured on individuals from the UK Biobank. We filtered the SNP genotype data based on minor allele frequency (≥ 0.01) and IMPUTE info score (≥ 0.3), leaving approximately 642,000 variants per trait. We used 42 environmental variables from the UK Biobank, similar to those used in previous GxE analyses of BMI in the UK Biobank ^28, 37^. After filtering on ancestry and relatedness, sub-setting down to individuals who had complete data across the phenotype, covariates and environmental factors we were left with approximately 280, 000 samples per trait. The sample, SNP and covariate processing and filtering applied is the same as that reported in the LEMMA paper^29^.

Table 1 shows the estimates and standard errors for SNP main effects 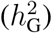 and GxE effects 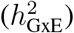 for GPLEMMA and LEMMA applied to the 4 traits. In all cases there is good agreements between the estimates from both methods.

**Table 1:** Comparison of GPLEMMA and LEMMA on 4 UK Biobank traits. Heritability estimates obtained using genotyped SNPs.

| Trait | $h_G^2$ (s.e) | | $h_{GxE}^2$ (s.e) | |
| --- | --- | --- | --- | --- |
|  | GPLEMMA | LEMMA | GPLEMMA | LEMMA |
| log BMI | 0.256 (0.078) | 0.259 (0.069) | 0.074 (0.008) | 0.071 (0.009) |
| PP | 0.230 (0.042) | 0.233 (0.039) | 0.063 (0.007) | 0.075 (0.018) |
| SBP | 0.237 (0.057) | 0.240 (0.053) | 0.036 (0.003) | 0.033 (0.003) |
| DBP | 0.273 (0.037) | 0.277 (0.034) | 0.021 (0.003) | 0.014 (0.001) |

## Discussion

Primarily this paper develops a novel randomized Haseman-Elston non-linear regression approach for modelling GxE interactions in large genetic studies with multiple environmental variables. This approach estimates GxE heritability at the same time as estimating the linear combination of environmental variables (called an ES) that underly that heritability. This general idea was pioneered in our previous approach LEMMA ^29^ which used a whole-genome regression approach to learn the ES, and this was then used in a randomized Haseman-Elston approach to estimate GxE heritability. The GPLEMMA approach introduced in this paper does not need that first whole-genome regression step, and this leads to substantial computational savings. The model underlying GPLEMMA is very similar to that in LEMMA, but implicity assumes a Gaussian distribution for main SNP effects and GxE effects at each SNP.

We compared GPLEMMA to a simpler approach, which we called MEMMA, that estimates GxE heritability of each environmental variable in a joint model, but does not attempt to find the best linear combination of them. We found that estimates of GxE heritability from MEMMA had higher variance than estimates from LEMMA and GPLEMMA, suggesting that the usefulness of MEMMA might be limited. Results from LEMMA and GPLEMMA were very similar, both in terms of estimating the ES and GxE heritability. The primary advantage of GPLEMMA over LEMMA is in computational complexity, as the empirical complexity of GPLEMMA appeared to be linear in sample size whereas LEMMA was shown to be super-linear ^29^.

In the future it may also be interesting to explore the idea of further partitioning variance using multiple orthogonal linear combinations of environmental variables. This could be expressed using the model

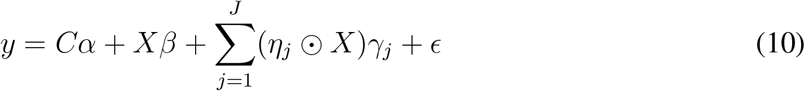

where *η_j_* = *E_wj_* is an N-vector, *w_j_* is an L-vector and 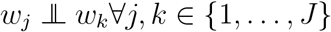.

LEMMA is also able to perform single SNP hypothesis testing whereas GPLEMMA (currently) does not. The linear weighting parameter *w* from GPLEMMA could be used to initialize LEMMA, or the estimated ES could be used as a single environmental variable in LEMMA. Exploring these, and other, approaches is future work.

## Acknowledgements

Computation used the Oxford Biomedical Research Computing (BMRC) facility, a joint development between the Wellcome Centre for Human Genetics and the Big Data Institute supported by Health Data Research UK and the NIHR Oxford Biomedical Research Centre. Financial support was provided by the Wellcome Trust Core Award Grant Number 203141/Z/16/Z. The views expressed are those of the author(s) and not necessarily those of the NHS, the NIHR or the Department of Health. We are grateful to Sriram Sankararaman for discussions about the RHE approach during the 2019 CGSI at UCLA.

## Author contributions

J.M. and M.K. conceived the ideas for the model and methods development. M.K. conducted all analyses and developed the software that implemented the methods with guidance from J.M. M.K. and J.M. wrote the manuscript.

## Footnotes

1 A parallel algorithm has perfect strong scaling if the runtime on T processors is linear in 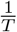, including communication costs.

## Notes

### Competing Interest Statement

The authors have declared no competing interest.

